# CoAtGIN: Marrying Convolution and Attention for Graph-based Molecule Property Prediction

**DOI:** 10.1101/2022.08.26.505499

**Authors:** Xuan Zhang, Cheng Chen, Zhaoxu Meng, Zhenghe Yang, Haitao Jiang, Xuefeng Cui

## Abstract

Molecule property prediction based on computational strategy plays a key role in the process of drug discovery and design, such as DFT. Yet, these traditional methods are time-consuming and labour-intensive, which can’t satisfy the need of biomedicine. Thanks to the development of deep learning, there are many variants of Graph Neural Networks (GNN) for molecule representation learning. However, whether the existed well-perform graph-based methods have a number of parameters, or the light models can’t achieve good grades on various tasks. In order to manage the trade-off between efficiency and performance, we propose a novel model architecture, CoAtGIN, using both Convolution and Attention. On the local level, k-hop convolution is designed to capture long-range neighbour information. On the global level, besides using the virtual node to pass identical messages, we utilize linear attention to aggregate global graph representation according to the importance of each node and edge. In the recent OGB Large-Scale Benchmark, CoAtGIN achieves the 0.0933 Mean Absolute Error (MAE) on the large-scale dataset PCQM4Mv2 with only 5.6 M model parameters. Moreover, using the linear attention block improves the performance, which helps to capture the global representation.

## I. Introduction

Throughout the human history of fighting diseases, small molecule drugs have played a vital role as the most reliable approach [1–4]. However, small molecule drug research is confronted with many challenges. It is estimated that 5,000-10,000 molecules need to be identified and validated in the discovery phase for one novel drug[5]. Considering the long and laborious period, researchers have summarized various guidelines to accelerate the drug discovery process based on drug-like character. For instance, the absorption, distribution, metabolism, excretion, and toxicity (ADMET) are highly related to the acid/base properties [6]. Moreover, the HUMO-LUMO gap, a quantum chemical property, might influence reactivity, photoexcitation, and charge transport [7]. Considering thousands of candidates, it is time-consuming and labour-intensive to do chemical or biological experiments for each compound.

With the development of chemistry and computer science, research has shown that the Force Field method can be employed to estimate some properties of molecules with high precision. For example, it is practicable to predict the force within and between molecules. Besides, Density Function theory (DFT) has become a valuable and helpful tool in many scenarios such as physics, chemistry, and materials [8]. This method contributes to drug discovery by precisely calculating various molecular properties such as the shape of molecules, reactivity, and energy gap [9]. Despite the superior success of DFT, the running time of the method is still costly for thousands of candidates, especially since it takes hours to calculate one single molecule’s properties.

Recently, artificial intelligence significantly contributed to the development of bio-pharmaceuticals. Several deep learning frameworks have been developed and performed well in the application of small molecule drug discovery [10], such as transformer-based BERT[11] and GPT[12]. The small molecule is generally represented using SMILES (simplified molecular input line specification) format. Thus, several works employed these well-verified pre-train language models to predict the molecule property. For example, SMILES-BERT [13] directly applied the BERT-style training strategy to the SMILES sequence using masked language learning. To overcome data scarcity, Chem-BERTa [14] utilized the multiple equivalent SMILES as input. Furthermore, TokenGT [15] used vanilla Transformer [16] to process molecule graph by treating atoms and bonds as independent tokens.

String-based representations mostly focus on sequence features, which are hard to capture the important topology information of molecules. If we regard atoms and bonds as nodes and edges of graph, using graph neural network (GNN) can help to obtain topology structure information, such as GIN [17], GAT [18], GIN-Virtual [19], Deeper-GCN [20]. These graph-based methods propagate messages of each node to its neighbours, and message-passing architecture can represent the molecule topology knowledge. However, the deeper and wider GNN-based models required more GPU memory. For example, the Edge-augmented Graph Transformer (EGT) employed a vanilla transformer to process node embeddings and introduced edge channels for each pair of nodes. This novel graph architecture has many model parameters (Fig. 1).

**Fig. 1.**
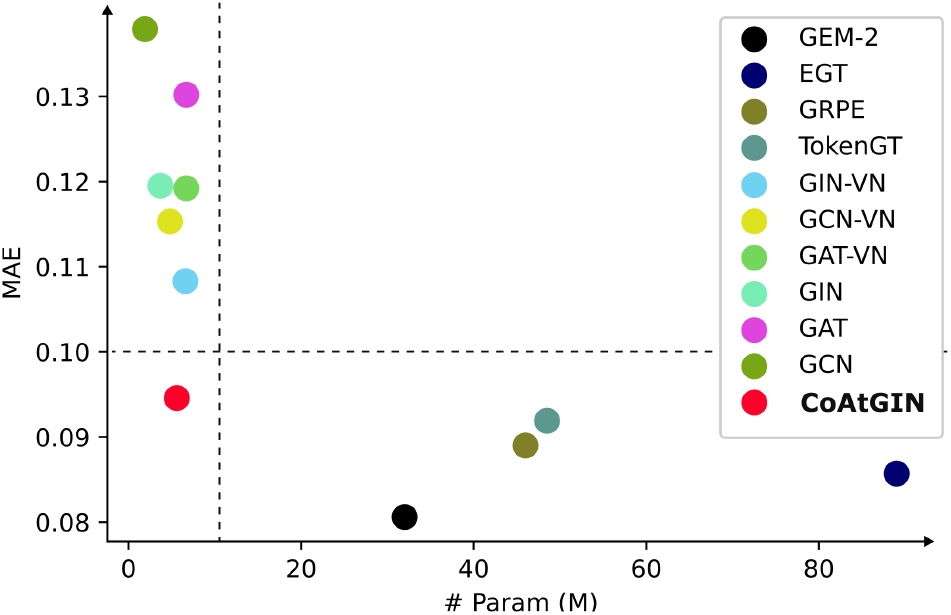
Models’ performance on the Open Graph Benchmark (OGB). The nodes in the upper-left area represent the models which perform not well enough but with fewer parameters. And the lower right area contains the well-performing model with enormous parameters. As the intersection area, CoAtGIN achieves a good grade while costs only a few parameters.

From the above discussion, we can observe that (a) String-based methods failed to encode the important topology information directly. GNN architecture is very suitable for processing small molecule drug data. (b) Virtual node passes identical messages to each node, which lacks the ability of global representation. (c) The scale of these methods keeps increasing to improve prediction accuracy (Fig. 1), which raises the challenge to manage the trade-off between efficiency and performance. Inspired by these observations, we proposed a novel graph-based architecture called CoAtGIN, the main contribution can be summarized as follows:

1. We present the k-hop convolution in a graph convolution network for faster message aggregation within one iteration.
2. We present a new way to accomplish global message passing through the graph using the linear transformer.
3. As shown in fig. 1, we develop a lightweight but well-performed model named CoAtGIN, which has only 5.2M parameters and achieves 0.0933 Mean Absolute Error (MAE) in the regression task of a large-scale graph dataset (PCQM4Mv2).

## II. Methods

As shown in Figure 2, CoAtGIN is composed of *L* layers. Each layer takes the *Node Embedding (NE)* and *Graph Embedding (GE)* as input, and then these two embeddings will be updated for layerwise iteration. Using the embedding block provided by [21], we initialize the *Node Embedding* as the atom type of each node. And the *Graph Embeddings* are set to zeros before the first layer.

**Fig. 2.**
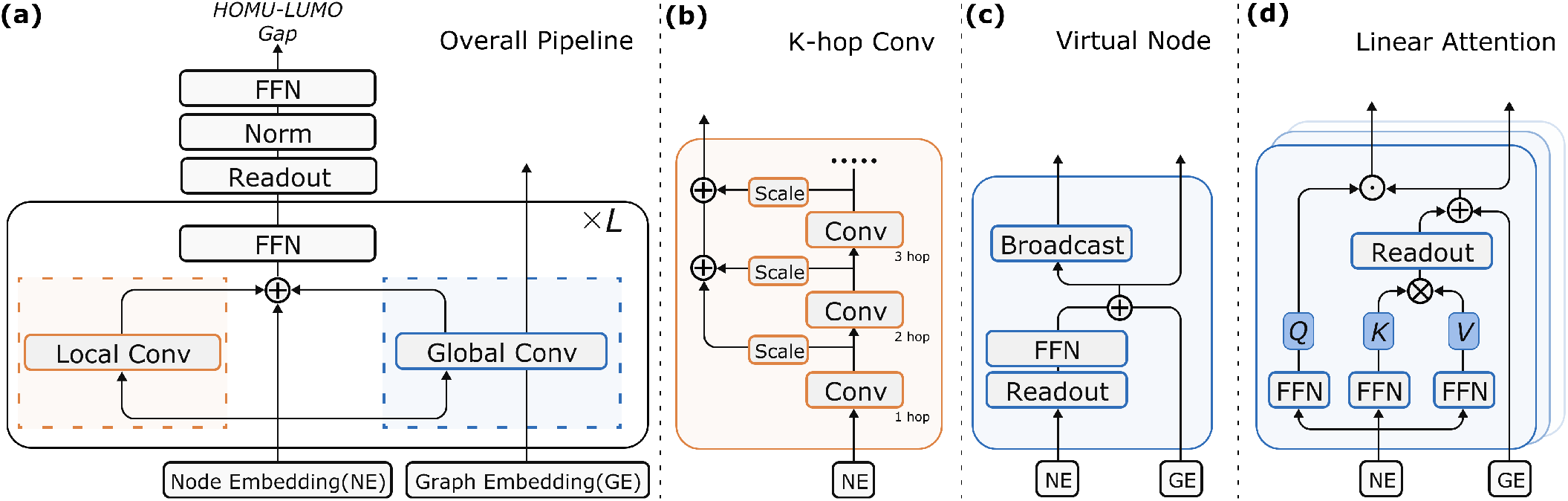
**(a)** The overall pipeline of the model for predicting the HUMO-LUMO gap, which consists of *L* repeated layers. Each layer can be broadly divided into two blocks, separately *Local Convolution* and *Global Convolution*, and classical residuals are used here. **(b)** *Local Convolution block*. The novel graph *local convolution* block named *k-hop convolution* repeatedly aggregated the neighbours’ information *k* times so that the *k*-hop nodes can be interacted in each iteration. **(c)** *Global Convolution block*. Illustration of *virtual node*, which reads from and writes to every node in the original graph separately using reduce and broadcast operations. GE (Graph Embedding) information from the last layer involves in the broadcast operation with the output of the feed-forward layer and the result also be used as newer GE. **(d)** *Global Convolution block*. Like the most linear transformer, the *K, V* matrix combines first and Q matrix do dot product with reduced global *KV*.

Although GIN-VN [19] wants to alleviate the lagging message passing problem between distant node pairs by adding the global virtual node, the broadcast message is identical to all the nodes. To solve this problem, section II-B shows shat we additionally apply the linear transformer as the *global convolution* block, which is the extension of GIN-VN. Besides taking both the linear attention and virtual node as *global convolution* block, we modify the traditional aggregation operation in *local convolution* block to efficiently capture the long-range neighbours’ features, which is shown in section II-C.

### A. Preliminaries

The Graph Isomorphism Network (GIN) [17] architecture was proposed by Xu *et al*. as a simple but powerful approach for the learning of graphs. It is illustrated and proved that GIN could effectively distinguish different graph representations. Moreover, the discriminative power of GIN is as strong as the 1-Weisfeiler-Lehman(1-WL) isomorphism test. Using the multi-layer perceptions (MLPs) to model the multiset functions, the propagation progress of GIN can be described as,

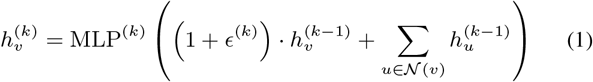

where 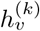 is the feature vector of node *v* at the *k*-th layer, MLP^(*k*)^ represents the *k*-th layer’s the multi-layer perceptions, *ϵ*^(*k*)^ denotes a learnable parameter, and *𝒩* (*v*) is the node set that contains all the neighbors of node *v*.

Nevertheless, the aggregation function shows that only neighbours can query and pass messages in the GIN and traditional graph network. Intuitively, the message through a distant node pair will be noisy and slow because the feature of one takes multiple iterations/layers to reach another. Therefore, Gilmer *et al*. [19] came up with a variant, GIN-VIRTUAL, which chose to add a virtual node to the graph with a particular edge type, and the node connects to all other nodes. Each original node in the graph reads and writes to this node in every step of message passing. Adding the virtual node has shown that the method works effectively across a wide range of graph-level prediction datasets [21].

However, as shown in Fig 2 (c), the virtual node of GIN-VN takes the READOUT as the graph representation and sends the duplicate processed message to every node in the original graph. Although Xu *et al*. [17] has proved that taking SUM as the graph level READOUT function can achieve good discriminative power. But each node in the graph also needs to accept different feature representations to optimize its representation.

### B. Global Convolution

Besides adding the virtual node, attention mechanism is another way to read and send information across the whole graph. Unlike adding the virtual node, the aggregated feature returned by the attention block differs from node to node. We apply the linear transformer to the global convolution block so that the diversity of the node can be guaranteed. Furthermore, we reserve the virtual node to enable the network to have a good ability both to distinguish different graphs and to gain a more expressive node representation.

#### 1) Virtual Node

For the *virtual node* block, the main pipeline maintains the same as GIN-VN. As shown in Figure 2 (c), the node embedding will be reduced the *Node Embedding (NE)* by the READOUT function (sum operation). All the node embedding will be added up to form the summarized information. The aggregated node feature will pass the feed-forward network to get the interpretation of the whole network, which we called *Virtual Node Graph Embedding (VGE)*

Added with the older *VGE*, the embedding enables updating the *Graph Embedding* of the virtual node and broadcasting the graph embedding for original nodes in the graph. It is notable that the broadcasted embeddings are identical for every node. However, different atoms play different roles in all kinds of chemical properties, which requires the network to feedback different information. Meanwhile, the extraordinary distinguish property should be reserved for identifying different molecules.

#### 2) Linear Transformer

To accomplish this task, the attention mechanism can be used here to afford the responsibility of calculating node-wise graph embedding. The result of the transformer is graph embedding driven by nodes, which we called *Attention Graph Embedding (AGE)*. By this way, both the node-wise and union representation graph embedding can be obtained in each iteration.

But the time complexity of the vanilla transformer is related to the squared number of nodes. In the desire of an efficient and lightweight model, we choose to use the CosFormer[22] to realize the attention mechanism for remedying the expensive computation cost,

On the basis of the CosFormer, we add the same READ-OUT function in the *virtual node* block (sum operation) for aggregating the global *KV*. Besides using the aggregated *KV* to update the *Graph Embedding* for linear attention, the matrix is also used to respond to the node-wise *Q*.

After the *linear attention* and *virtual block* generating the *AGE* and *VGE*, the two embeddings are added together as the output of *global convolution* block.

### C. K-Hop Convolution

*Linear attention* and *virtual node* block are both the way of adding new message passing methods throughout the network. In terms of the vanilla node aggregation operation as shown in eq. (1), only the features within 1-hop distance can be captured.

By repeating the propagation *k* times, we can receive the at most *k*-hop distance information. It makes it possible to interact with the long-range node without the use of a deep Graph Neural Network. And all the aggregation result will be scaled by a learning layer so that the node can capture import neighbours and bonds.

The ablation study in section III-C shows that the model benefits a lot due to the k-hop convolution.

## III. Experiments

In this paper, we evaluate the performance of the proposed architecture CoAtGIN using PCQM4Mv2 dataset, which contains 3.8 million molecular graphs. The supervised setting is a quantum-chemical regression task, which predicts the DFT-calculated HOMO-LUMO energy gap. The HOMO-LUMO energy gap is a human-defined important metric in quantum chemistry, which is strongly associated with many molecule properties such as polarizability and activation energy. Moreover, Mean Absolute Error(MAE) is used to evaluate the benchmark accuracy, and the lower value indicates a better model performance.

Besides, we instigate the representation of CoAtGIN by visualizing the graph embedding. The result indicates that the graph representation can describe the molecule well, so embedding visualization is continuous.

At last, we explore whether the modules in our architecture make sense in this quantum-chemical regression task. Based on the observation, further exploration might be obtained by the experiment.

### A. Comparison to the state of the art methods

As mentioned before, the CoAtGIN model is evaluated in the Large Scale Challenge (LSC) benchmark [21] PCQM4Mv2 dataset. There are many fancy models using the benchmark as their evaluation metrics. As shown in table I, which includes the state-of-the-art methods and their corresponding asymptotics, number of parameters, and MAE result. The models here can be broadly divided into two types by the model’s number of parameters.

**TABLE I.** Result on PCQM4Mv2 large-scale graph dataset.

| Method | Asymptotics | # Param | Validate MAE ↓ |
| --- | --- | --- | --- |
| GCN | $\mathcal{O}(n + m)$ | 2.0M | 0.1379 |
| GAT | $\mathcal{O}(n + m)$ | 6.7M | 0.1302 |
| GIN | $\mathcal{O}(n + m)$ | 3.8M | 0.1195 |
| GAT-VN | $\mathcal{O}(n + m)$ | 6.7M | 0.1192 |
| GCN-VN | $\mathcal{O}(n + m)$ | 4.9M | 0.1153 |
| GIN-VN | $\mathcal{O}(n + m)$ | 6.7M | 0.1083 |
| **CoAtGIN** | **$\mathcal{O}(n + m)$** | **5.2M** | **0.0933** |
| TokenGT | $\mathcal{O}((n + m)^2)$ | 48.5M | 0.0910 |
| GRPE | $\mathcal{O}(n^2)$ | 46.2M | 0.0890 |
| EGT | $\mathcal{O}(n^2)$ | 89.3M | 0.0869 |
| Graphormer | $\mathcal{O}(n^2)$ | 48.3M | 0.0864 |

The traditional Graph Neural Network like Graph Convolutional Network (GCN) [23], Graph Attention Network (GAT) [18], Graph Isomorphism Network (GIN) [17] and their variants (GCN-VN, GAT-VN, and GIN-VN) have only a few parameters but good performances with about 0.1 MAE. However, to achieve the better performance, TokenGT [24], GRPE [25], EGT [26] and Graphormer [27] adopts plenty of fancy modules and architecture. Although the performance does improve, the number of parameters increases severely. Even the slightest TokenGT [24], whose number of parameters is 7.2 times larger than the GIN-VN model.

Although the scale of CoAtGIN is only 5.2M, the model achieves the 0.945 MAE on PCQM4Mv2 dataset which is close to the result of TokenGT. Besides, the asymptotic columns show that the time complexity of CoAtGIN is one of the lowest models in the benchmark. In the words, this experiment shows that CoAtGIN is a lightweight model which can achieve fast inference speed without much loss of performance.

### B. Investigation of CoAtGIN representation

We analyse the molecule representation learned by CoAtGIN using T-distributed Stochastic Neighbor Embedding(t-SNE) [28] method and the scikit-learn [29] package. t-SNE can map high-dimensional space into two-dimensional space, showing the embedding information intuitively. We random sample 2000 molecules from PCQM4Mv2 dataset to illustrate what the CoAtGIN learns. From Fig. 3, the shorter the distance of the molecule representations, the closer the quantum-chemical property value is. In other words, similar 2D representations share close molecule properties. For example, the light blue indicates a low HOMU-LUMO gap value, and we can see that these blue dots are clustered together. More interestingly, the 2D embedding representation is continuous in the t-SNE space, which is high relative to the continuous HOMU-LUMO gap value (regression value). This indicates that molecules with similar properties have close features using CoAtGIN for embedding. If we regard CoAtGIN as an effective pre-train strategy, the proposed method will likely perform better in downstream tasks.

**Fig. 3.**
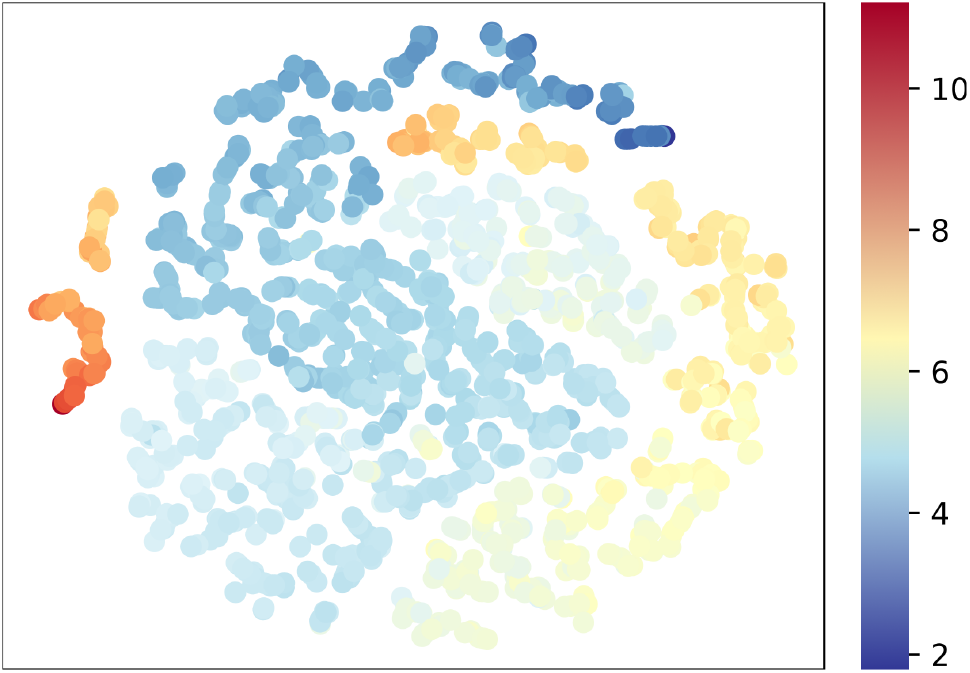
T-SNE visualization of the aggregated nodes’ embedding. The color indicates the ground truth of the molecules’ HOMO-LUMO gap. The dark blue represents the lowest energy gap and the red color represents the highest value.

### C. Ablation experiments

To explore the contribution of each module illustrated in Fig. 2 and section II, we evaluate these modules using an ablation study. There are three blockes mentioned in section II: *k-hop conv, virtual node*, and *linear attention*. And the ablation experiment is detailed in table II.

**TABLE II.** Ablation study of the modules in CoAtGIN.

| K-Hop Convolution (k = ) | Virtual Node | Linear Attention | # Param | Validate MAE ↓ |
| --- | --- | --- | --- | --- |
| 1 | — | — | 1.8M | 0.1084 |
| 2 | — | — | 2.5M | 0.1012 |
| 3 | — | — | 2.5M | 0.1002 |
| 3 | ✓ | — | 3.8M | 0.0945 |
| 3 | — | ✓ | 3.9M | 0.0941 |
| 3 | ✓ | ✓ | 5.2M | **0.0933** |

Firstly, we illustrate the relationship between the performance and the *k* value selection of *k-hop convolution* block. From the table II, we can see that the model performance positively correlates with the hop convolution. A higher *k* value usually brings better performance which means that further node information is helpful to capture better local and topology information. Considering the trade-off between efficiency and performance, larger *k* values were not validated because the 3-hop neighborhood contains most of the necessary bioinformatic information like torsion angles. Larger *k* values will lead to the duplicated information problem and make the model larger. Based on this observation, we finally choose *k* = 3 in the *k-hop convolution* block considering both the performance and efficiency.

And then we explore the usage of *linear attention* and *virutal block* as shown in the lower part of table II. Table II indicates that virtual node and linear attention both play a key role in CoAtGIN. Using any of the *linear attention* and *virutal block* in CoAtGIN makes the regression value much more precise. The combination of these two modules further improves the performance. It is convincing that the *linear attention* and *virutal block* are both necessary for the network, and learning features from different modules might improve the other.

## IV. Conclusion

In summary, we propose a novel graph-based architecture, CoAtGIN, to obtain node embedding and graph embedding. Experiments indicate that CoAtGIN achieves competitive performance with relatively high efficiency in the OGB Large-Scale Challenge. The key modules of CoAtGIN include k-hop conv, virtual node, and linear attention. The k-hop conv can capture better local typology structure information. And the linear transformer is designed to remedy the problem of identical messages from the virtual node. Using both convolution and attention provides a novel way to predict molecule properties.

In the future, we will continue to explore the strategies for improving CoAtGIN. The rotation angle plays an essential role in the structure and function. Our next direction is to design an effective block to capture the geometric properties of molecules. We also want to understand why CoAtGIN performs better from the biological perspective. In addition, our proposed CoAtGIN can effectively process graph data, so this model could be a good fit to solve protein structure-related problems. Because we regard residues as nodes, the contact between residues is defined as edge. More exciting applications of CoAtGIN for protein complex prediction and design will be explored.

